# Towards precision ecology: Relationships of multiple sampling methods quantifying abundance for comparisons among studies

**DOI:** 10.1101/2021.09.14.460401

**Authors:** Benjamin D. Hoffmann, Magen Pettit

## Abstract

Because different sampling techniques will provide different abundance values, it is currently difficult to compare results among many studies to form holistic understandings of how abundance influences ant ecology. Using three sampling methods in the same location we found pitfall traps best confirmed *A. gracilipes* presence recording the fewest zero values (9.1%), card counts were the least reliable (67.1%), and tuna lures were intermediate (30.1%). The abundance of *A. gracilipes* from card counts ranged from 0 to 20, in pitfall traps from 0 to 325, and the full range of tuna lure abundance scores (0-7) were sampled. We then determined the relationships between these three standard ant sampling techniques for the abundance of yellow crazy ant *Anoplolepis gracilipes*. Irrespective of the data transformation method, the strongest relationship was between pitfall traps and tuna lures, and the least strong was between pitfall traps and card counts. We then demonstrate the utility of this knowledge by analysing *A. gracilipes* abundance reported within published literature to show where the populations in those studies sit on an abundance spectrum. We also comment on insights into the relative utility of the three methods we used to determine *A. gracilipes* abundance among populations of varying abundance. Pitfall traps was the most reliable method to determine if the species was present at the sample level. Tuna lures were predominantly reliable for quantifying the presence of workers, but were limited by the number of workers that can gather around a spoonful of tuna. Card counts were the quickest method, but were seemingly only useful when *A. gracilipes* abundance is not low. Finally we discuss how environmental and biological variation needs to be accounted for in future studies to better standardise sampling protocols to help progress ecology as a precision science.

## INTRODUCTION

There is a myriad of ways to sample ant abundance [1,2]. For clarity, all use of the word abundance in this paper refers solely to “momentary above-ground worker abundance” from 30 second counts to 24 hour counts, not overall abundance which would incorporate non-foraging workers. Although there will likely be a positive relationship between forager abundance with overall population levels and worker density [3], these relationships and other potential metrics are not considered further. To quantify abundance, the sampling method of choice for any study will depend upon many factors including effort, desired sampling completeness, the science questions, the foraging biology of the target species, and the data requirements of proposed analyses. But because different sampling techniques will provide different abundance values, it is currently difficult to compare results among many studies to form holistic understandings of how abundance influences ant ecology, and largely impossible for quantified comparisons to give predictive knowledge. This issue is most prominent when there is little complementarity between both sampling technique and the environments (e.g. card counts in a rainforest vs pitfall traps in an open environment).

Most research investigating comparability of ant sampling methods focuses on sampling entire faunas (species richness) and has investigated variations of the same type of method such as comparing different pitfall trap diameters [4] or preservatives [5,6], as well as comparing data collected using different sampling methods [8,9,10,11]. Assessments focusing on individual species appear to be largely restricted to matrix preferences of invasive species for toxic bait attractancy [e.g. 12,13]. For all such work, as far as we are aware, there has never been a comparative study that has produced quantitative relationships of the outcomes of using different methods that can be used to compare dissimilar data among other studies.

Resolving the issue of non-complementarity of data among studies will be one of the most productive ways to accelerate knowledge and address key science questions. One of the best examples of the need to address this data issue is the attempts to gain a predictive understanding the impacts of invasive species. Impacts of biological invasions are density dependent [14,15,16], so having data that can be precisely compared among studies sampling the target invader is critical if we are to obtain truly predictable and quantifiable understandings of the effects of invasive species. To some extent we can make comparisons and broad generalisations when sampling and environments are similar, or even when sampling is consistent among very different environments, but even this output remains far from quantifiable.

Here, we attempt to partly resolve this issue by determining the relationships between three standard ant sampling techniques for the abundance of the invasive yellow crazy ant *Anoplolepis gracilipes*. We then demonstrate the utility of this knowledge by analysing *A. gracilipes* abundance reported within published literature to show where the populations in those studies sit on an abundance spectrum. We also comment on insights into the relative utility of the three methods we used to determine *A. gracilipes* abundance among populations of varying abundance.

## MATERIAL AND METHODS

### Study area

The study was conducted within northeast Arnhem Land (12°11’S, 136°46’E) in Australia’s Northern Territory. The regional climate is tropical monsoonal with high temperatures (17-33C) throughout the year and an annual rainfall of approximately 1200 mm falling predominantly during the summer wet season. The landscape is predominantly savanna woodland dominated by *Eucalyptus tetrodonta* (height and canopy cover approximately 15 m and 20% respectively), with an understorey to three meters predominantly of Acacias and grasses [17]. The weather was consistent throughout the sample period, being dry and sunny (temperature range 21-32C) and all sites were sampled simultaneously, therefore we do not consider that weather variation influenced the results.

### Sampling

We quantified *A. gracilipes* abundance at three sites, being three spatially discrete *A. gracilipes* populations with visually different active ant abundance. This species occurs throughout the region in discrete populations [18], and all populations were familiar to the authors as they had been recently surveyed in preparation for eradication [19]. Sampling was conducted along transects, with 11 sampling locations along each transect, and both transects and sampling locations spaced 10 m apart. There were 13 transects at two sites with visually low and medium-level *A. gracilipes* abundance, and 15 transects at the third site with visually high abundance. Within a site a single person conducted sampling on each transect and a different person was used for each transect. Sampling was conducted using three methods at all three sites, being card counts, tuna lures and pitfall traps. A single sample of each method was used at each sample location in three separate sampling occasions. Card sampling was conducted first at two sites in the late afternoon (after 4 pm) of 18 September 2009. The following day tuna lures were used at the same two sites at exactly the same time of day. Pitfall traps were set the following day and operated for 24 hours. The same sampling order and timing was conducted at the third site commencing 21 September 2009.

Cards were 10 cm x 10 cm laminated paper. At each sampling location a card was placed on the ground with the edges in contact with substrate as far as possible to allow easy access for the ants to walk onto the card. The card was observed for 30 seconds with the number of *A. gracilipes* workers that walked over the square recorded.

Tuna lures were a teaspoon amount of canned tuna (Home brand Seafood Basket cat food containing tuna). At each sampling location a lure was placed directly onto the ground and *A. gracilipes* abundance at and within 1cm of each lure was scored after 20 minutes according to the following scale that has been applied to many publications containing ant abundance datasets: 0 = no ants; 1 = 1 ant; 2 = 2-5 ants; 3 = 6-10 ants; 4 = 11-20 ants; 5 = 21-50 ants; 6 = 50-100 ants; and 7 = >100 ants. This scale is used because of the impracticality of accurately counting large numbers of ants at a lure.

Pitfall traps were plastic containers with an internal diameter of 65 mm, one third filled with ethylene glycol as a preservative. A single pitfall trap was set at each sample location, operated for 24 hours, collected, and *A. gracilipes* abundance was counted back in a laboratory. Because of logistics constraints only 13 transects were sampled using pitfall traps, with these transects chosen to cover the full abundance variation found prior by card counts and tuna lures. Six, four and three transects were sampled at the three sites respectively (Figure 1).

**Figure 1.**
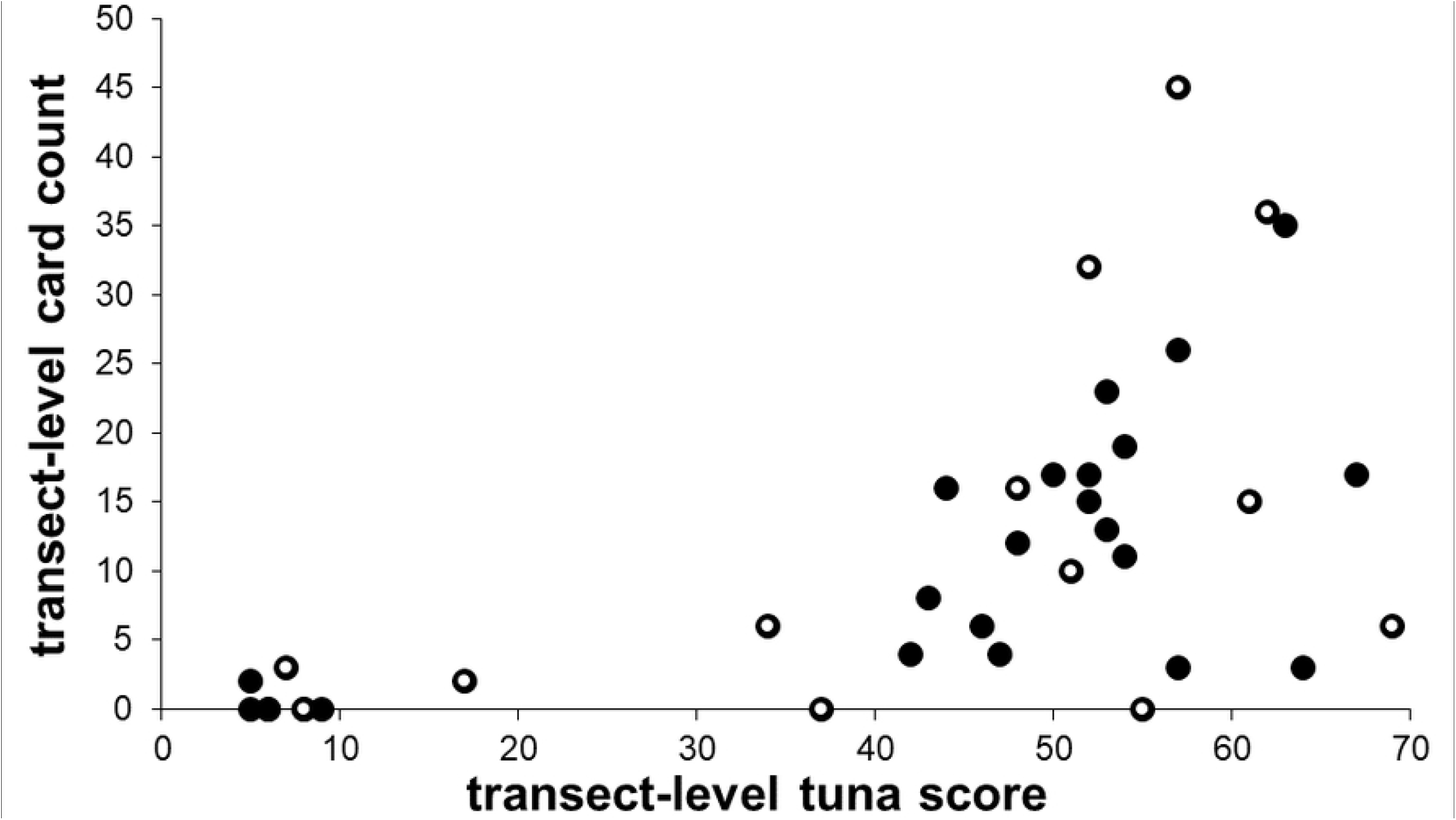
Transect-level combined counts of tuna lures and card counts showing transects that were (white circles) and weren’t (black circles) sampled with pitfall traps.

### Analysis

Four of the transects in the low-abundance site obtained no *A. gracilipes* in both card counts and tuna lures, and were also not subsequently sampled using pitfall traps, so these transects were excluded from analyses, predominantly because we could not confirm if *A. gracilipes* were definitely present or not when the data were analysed. First, the scaled data of the tuna lures was transformed to give an actual abundance value, being the value or mid-range value of each abundance category, being: 0 = 0; 1 = 1; 2 = 3; 3 = 8; 4 = 15.5; 5 = 35.5; 6 = 75.5; and 7 = 125. Next, the abundance data for each sampling location were transformed to reduce the spatial scatter of the data, and was done so two ways to compare the two most extreme mathematical possibilities. First was a log(x+1) transformation on data first summed at the transect-level (data of 11 samples first combined and then transformed) and second was sample-level data first transformed and then summed to transect-level. The data were then compared first pairwise in 2D scatterplots fitted with linear Pearson correlations, then together in 3D scatterplots.

To demonstrate the utility of these correlations we then compared ant abundances among published studies. First, we identified 12 publications that sampled yellow crazy ants using one of these methods and also provided data. Although most studies contained numerous sites with varying levels of crazy ant abundance, we only used the data from the sampling unit with the highest abundance. Where possible we obtained sample-level data, but in most instances all that was available was an average per sample or a total of summed samples (e.g. 5 pitfall traps summed as a plot), in which case we averaged the data to obtain an abundance value per sample, and if needed standardised the data to the methods used here (e.g. halving the data of the 1-minute card count in [20] to match the 30 second card count used here). Because traps were used the most (5 out of 11 studies), we used the correlation equations to calculate abundance in traps for the other six studies. To ensure sample number was the same among studies all calculations were made using 11 samples as the sample number to make data comparable with this study. This was done for both data transformation methods. These trap data were then used to calculate all remaining data of the other two sample types for each study using the same methodologies, but by re-arranging the y=ax+b equations to x=(y-b)/a. The data were then visually compared graphically.

## RESULTS

At the sample level, where all three sampling methods were used, pitfall traps best confirmed *A. gracilipes* presence recording the fewest zero values (13; 9.1%), card counts were the least reliable (96; 67.1%), and tuna lures were intermediate (43; 30.1%). In the 13 instances where *A. gracilipes* workers were not present in a trap, no *A. gracilipes* were also recorded by both other methods in 12 instances, and a single worker was recorded at the tuna lure in the other instance, confirming that *A. gracilipes* was largely absent from these few locations. The abundance of *A. gracilipes* from card counts ranged from 0 to 20, in pitfall traps from 0 to 325, and the full range of tuna lure abundance scores (0-7) were sampled.

The raw sample-level data showed great variation (Figures 2a-c), but naturally this variation was greatly reduced after combinations of log transformation and pooling (Figures 2d-i). Irrespective of the data transformation method, the strongest relationship was between pitfall traps and tuna lures, and the least strong was between pitfall traps and card counts. The two transformation and pooling methodologies gave similar but distinctly different results. However, the 3D compilation of sample-level data that had undergone transformation prior to pooling provided a greater data spread (Figure 3). Notably two transects with intermediate *A. gracilipes* abundances that also had zero values on card counts look anomalous in the figures.

**Figure 2.**
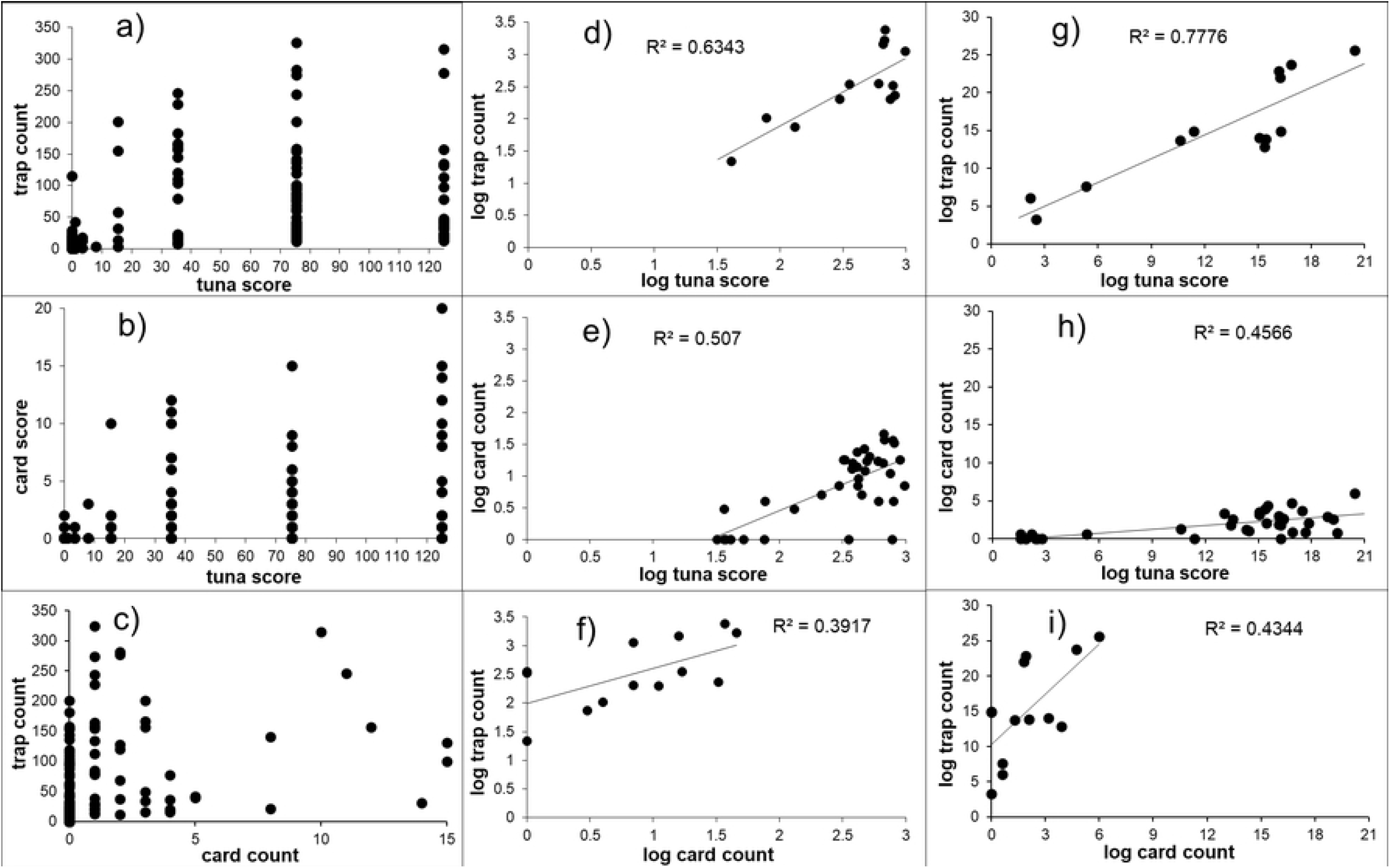
Pairwise relationships of the three sampling methods of card counts, pitfall trapping and tuna lures with raw sample-level data (a-c), transect-level data compiled from first pooling sample-level data then applying a log(x+1) transformation (d-f) and transect-level data compiled by pooling sample-level data that had first undergone a log(x+1) transformation (g-i).

**Figure 3.**
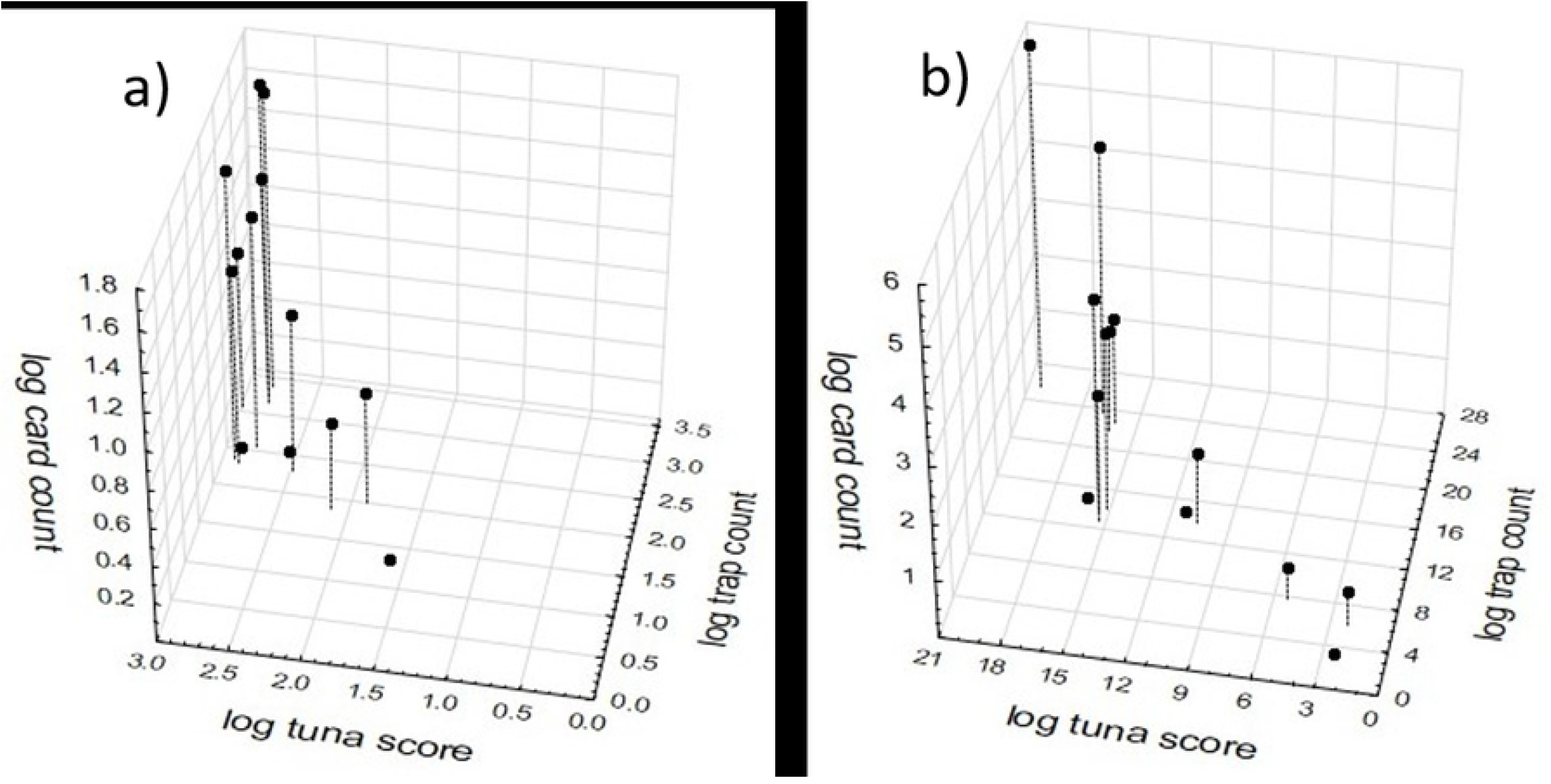
3D scatterplots of the relationship between *A. gracilipes* abundance determined by the three sampling methods of card counts, pitfall trapping and tuna lures using transect-level data compiled from first pooling sample-level data then applying a log(x+1) transformation (a) and transect-level data compiled by pooling sample-level data that had first undergone a log(x+1) transformation (b).

The standardisation calculations of pitfall trap abundance from published data using the two transformation and pooling methodologies (Figure 4) ordered the publications very similarly. But the two methods gave greatly different results for the two studies with greatest crazy ant abundance (studies 11 and 12) whose original data were based on card counts, with the method that used log transformation of already pooled data (Figure 4b) providing a seemingly artificially low abundance count. The 3D compilation of sample-level data that had undergone transformation prior to pooling largely maintained the order of the studies as determined by pitfall trap data, but with some clear nuances of lower abundances relative to the general pattern, predominantly (2 of 3 studies) associated with studies that whose data were based on card counts (studies 2 and 11) (Figure 5).

**Figure 4.**
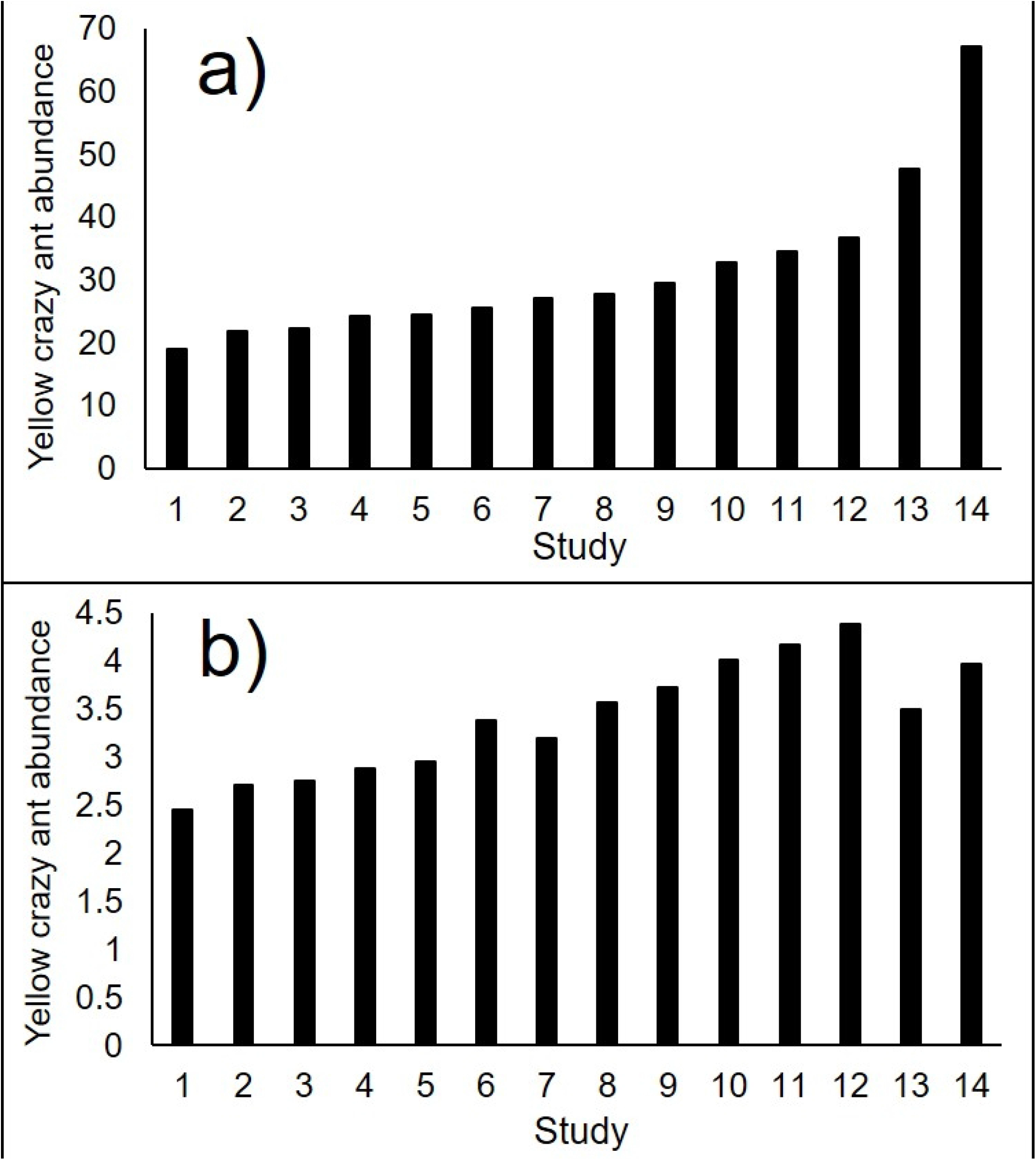
Standardised highest yellow crazy ant abundance reported within fourteen studies as calculated by the relationships between the three sampling methods of card counts, pitfall trapping and tuna lures using a) summed log(x+1)-transformed sample data, and b) log(x+1)-transformed pooled data. Studies are: 1) [30]; 2) [31] New Caledonia site; 3) [31] Australia site; 4) [20]; 5) [32]; 6) this study; 7) [33]; 8) [34] 9) [18]; 10) [35]; 11) [36]; 12) [37]; 13) [38]; and 14) [3].

**Figure 5.**
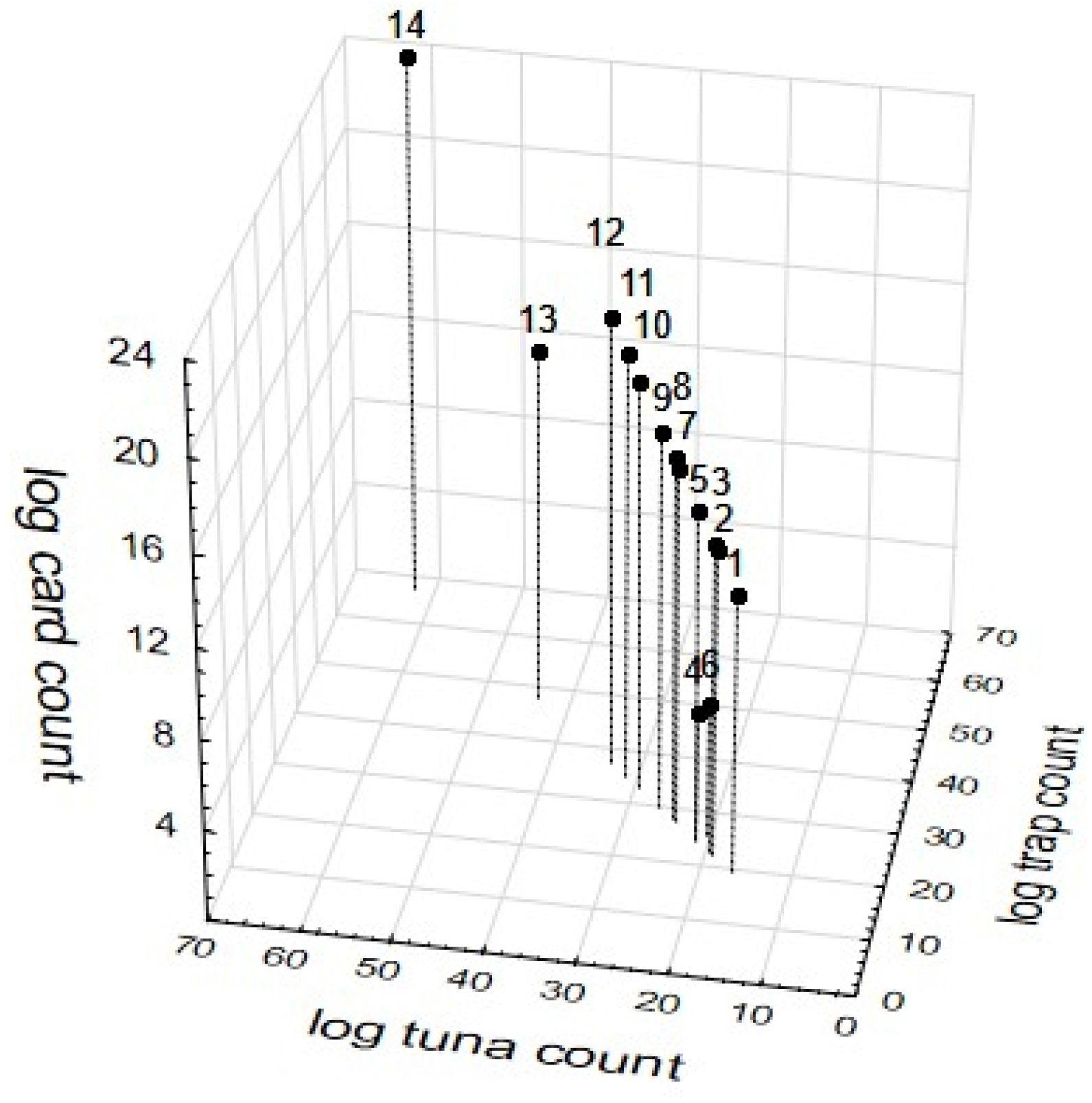
3D scatterplot of standardised highest yellow crazy ant abundance among fourteen studies as calculated by the relationships between the three sampling methods of card counts, pitfall trapping and tuna lures using sample-based log(x+1)-transformed pooled data. Studies are: 1) [30]; 2) [31] New Caledonia site; 3) [31] Australia site; 4) [20]; 5) [32]; 6) this study; 7) [33]; 8) [34] 9) [18]; 10) [35]; 11) [36]; 12) [37]; 13) [38]; and 14) [3].

## DISCUSSION

Despite great differences in sampling methodology, the abundance data obtained from the three sampling techniques attained reasonably strong relationships, but only after transformation and pooling. This outcome provided a strong basis by which abundance among studies using different sampling techniques could be calculated and compared. Here we used two types of transformations and found similar but distinct results, and ultimately of these two we recommend the use of the data that transforms samples before pooling. However, this does not preclude the use of other transformations that may prove to be better. Indeed, we have provided our raw data as Supplementary Material to encourage other researchers to explore other mathematical relationships, and perhaps other analytical techniques, that may prove to be superior to what we have found here.

Even with the relatively small gradient of *A. gracilipes* abundance assessed within this study, coupled with universal implications for their use, the three techniques had clearly different capacity and practicality for use. Pitfall traps had the lowest occurrence of absence data, so was the most reliable method to determine if the species was present at the sample level. The downside of this method though is that it is very time consuming and laborious and does not give results in real-time because the ants need to be counted back in a laboratory. Pitfall traps are also prone to prone to inflated abundance data if a trap is positioned close to a nest. Tuna lures were predominantly reliable for quantifying the presence of workers, but were limited by the number of workers that can gather around a spoonful of tuna, being not too many more than 100 for *A. gracilipes*, which can also be influenced by user-induced variation in lure size and shape. Therefore lures (at least of this size) would be less useful in areas with high and very high *A. gracilipes* abundance. Card counts were the quickest method, but were seemingly only useful when *A. gracilipes* abundance is not low. In the most extreme case, a zero count was obtained by a card, but the associated pitfall trap collected 201 workers. Notably cards are not limited by ant numbers like tuna lures are, and also potentially pitfall traps, and so are great to use when ant abundance is very high.

What remains unclear is how variation with the ant’s diurnal foraging cycle [21], prevailing environmental conditions, and annual population cycle would affect results and ultimately the mathematical relationships. Because pitfall traps are operated normally for 24 to 48 hours, sometimes up to a week, this method has the advantage that it is not so subject to short-term variations in environmental conditions, and captures forager abundance throughout full diurnal cycles. But ant abundance at lures, and especially card counts due to their “snapshot” assessment times” will be completely subject to momentary abundance at that point on the diurnal cycle coupled with effects of prevailing environmental conditions. So for both methods, different abundance values would likely be attained at different stages of the diurnal cycle and between when environmental conditions are ideal vs less ideal. Similarly, attendance at lures may also be influenced by the presence of other resources and dietary needs of the colony at the time relative to the composition of the lure. Indeed even just lure placement in sunny vs shaded locations is known to affect ant presence [22]. As a prediction for all epigeic ant species, we suspect that such variation issues will be greater in open environments more than closed environments due to the effects of solar insolation, and relatively greater in other locations where there are big changes in daily temperature such as on mainland systems relative to lowlands of island systems. Finally, all three methods would also give differing results between peak and low times of the ant’s annual population cycle.

The combined analysis of published work we presented here does not account for any of the variation issues detailed above, and so therefore should be interpreted with some caution and not seen as being definitive. Indeed even just the differences in the results between the two data transformations used demonstrates the lack of definitive nature of the analysis. Regardless, the calculations appear to confirm the claim that yellow crazy ant abundance quantified on Christmas island within [3] are the highest of all reported for yellow crazy ant, and probably remains the highest quantified for any ant species in the world.

Our study has advanced the potential for meta-analyses of *A. gracilipes* studies, by providing a mathematical way to calculate abundance that is standardised among studies that have quantified abundance using either pitfall traps, lures or cards to determine an abundance spectrum. But we have not yet been able to advance the utility of many studies that use other methods to quantify *A. gracilipes* abundance using methods like insecticide knockdown, standardised hand collections, foliage beating, sugar-soaked pads [23,24,25,26,27,28) and of course when abundance data are not used or reported (e.g. 29). It remains unclear how universal the relationships would be for other ant species. We suspect that these relationships would not be universal, but may be similar enough for ecologically similar species that have similar abundance, similar locomotion speeds, and similar foraging strategies. Ultimately, understanding and accounting for momentary, daily and seasonal variations in ant abundance, even avoiding these issues altogether within future studies perhaps through the use of standardised sampling protocols, is a great science challenge that overcoming will help progress ecology as a precision science.

## ACKNOWLEDGEMENTS

We thank Lori Lach, Peter Caley and Petra Kuhnert for conversations about analysing and interpreting the data. Lori Lach, Petr Klimes, and two anonymous reviewers provided comments on the draft manuscript.

